# BWGS: a R package for genomic selection and its application to a wheat breeding programme

**DOI:** 10.1101/763037

**Authors:** Gilles Charmet, Louis Gautier Tran, Jérôme Auzanneau, Renaud Rincent, Sophie Bouchet

**Affiliations:** INRA-UCA, UMR GDEC– 5 chemin de Beaulieu, 63000 Clermont-Ferrand, France; Agri-Obtentions, Ferme de Gauvilliers, 78660 Orsonville, France

## Abstract

We developed an integrated R library called BWGS to enable easy computation of Genomic Estimates of Breeding values (GEBV) for genomic selection. BWGS relies on existing R-libraries, all freely available from CRAN servers. The two main functions enable to run 1) replicated random cross validations within a training set of genotyped and phenotyped lines and 2) GEBV prediction, for a set of genotyped-only lines. Options are available for 1) missing data imputation, 2) markers and training set selection and 3) genomic prediction with 15 different methods, either parametric or semi-parametric.

The usefulness and efficiency of BWGS are illustrated using a population of wheat lines from a real breeding programme. Adjusted yield data from historical trials (highly unbalanced design) were used for testing the options of BWGS. On the whole, 760 candidate lines with adjusted phenotypes and genotypes for 47 839 robust SNP were used. With a simple desktop computer, we obtained results which compared with previously published results on wheat genomic selection. As predicted by the theory, factors that are most influencing predictive ability, for a given trait of moderate heritability, are the size of the training population and a minimum number of markers for capturing every QTL information. Missing data up to 40%, if randomly distributed, do not degrade predictive ability once imputed, and up to 80% randomly distributed missing data are still acceptable once imputed with Expectation-Maximization method of package rrBLUP. It is worth noticing that selecting markers that are most associated to the trait do improve predictive ability, compared with the whole set of markers, but only when marker selection is made on the whole population. When marker selection is made only on the sampled training set, this advantage nearly disappeared, since it was clearly due to overfitting. Few differences are observed between the 15 prediction models with this dataset. Although non-parametric methods that are supposed to capture non-additive effects have slightly better predictive accuracy, differences remain small. Finally, the GEBV from the 15 prediction models are all highly correlated to each other. These results are encouraging for an efficient use of genomic selection in applied breeding programmes and BWGS is a simple and powerful toolbox to apply in breeding programmes or training activities.

## Introduction

The use of molecular markers to provide selection criteria for quantitative traits was first proposed by [1]. They introduced a theory for optimizing weights to be given to each marker associated with a QTL, and they demonstrated that this index was at least as efficient as the phenotypic score for the genetic improvement of a population by truncation selection. This marker-assisted-selection (MAS) approach used only markers which had been previously associated with a QTL through statistical analysis. Therefore, the number of markers remained limited, and their effects were usually estimated by solving the linear model equations. The efficiency of MAS versus phenotypic selection is higher when the trait has a low heritability, the population size is large and the QTLs explain a large proportion of the trait variation. It was shown that the use of this marker index would facilitate early selection, bypassing a trait evaluation step and thereby shortening selection cycles [2]. Consequently the genetic gain per cycle would increase. However, QTL detection is often limited in practice by experimental power, particularly by the size of the studied population, and QTL with small effects often remain undetected. Therefore, this “missing heritability” can reduce the efficiency of MAS. Subsequent studies have shown that MAS efficiency is improved when more QTLs with small effects are included [3–4], and this implied relaxing the stringency threshold of significance to allow more true QTLs being detected, despite the risk of having some false positive ones being included. Extending this reasoning, it was proposed to include all markers in the selection index, thus bypassing the QTL identification step [5]. As the number of markers is generally larger than the number of phenotypic observations, classical fixed effects regression are intractable (n>>p problem). Therefore, [5] suggested using ridge regression models to overcome this over-parameterization problem. Soon after, [6] applied ridge and Bayesian regression models to animal populations for predicting breeding values, and called this approach Genomic Selection (GS). Marker effects are first estimated from the genotypic and phenotypic data in a training population. Then marker effects are used to predict breeding values in the target population with only genotypic data, and selections are based on these Genomic Estimates of Breeding Values (GEBV). This method has been used successfully for dairy cow breeding [7]. Indeed, in the case of dairy cow breeding (and particularly for bulls), the advantages of GS over classical breeding are obvious. 1) genotyping is much cheaper (a few 10s US$ for the 55K SNP bovine chip) than progeny testing (since bulls do not give milk), 2) genotyping can be done at birth time, while progeny testing requires >7-8 years (until bulls give daughters and daughters give milk). Therefore GS of dairy bulls allowed early selection on a larger population, thus leading to nearly doubling the genetic gain per unit of time while the costs of proving bulls were reduced by 92% [8].

To benefit from shorter cycles and increasing selection intensity, the genetic gain per cycle should be close to that expected from phenotypic selection (PS). This relative efficiency of GS vs PS thus relies on the ability of predicting the observed genetic value from the marker genotype. GEBV predictive ability is usually measured by the correlation between the predicted and observed values. Most reported studies on GS in plants have focused on measuring the accuracy of genomic predictions, usually assessed by cross-validation techniques [9–14].

In the native theory, all available markers should be used without prior selection, since the statistical models were supposed to cope with the big data problem and avoid over parametrization. However, with the advance of molecular biology, the number of markers, e.g. single nucleotide polymorphisms (SNP), can be extremely high (up to some millions), particularly when compared to a few hundreds or thousands phenotypic observations. In this case, it seems reasonable to limit the number of markers to avoid too high over parametrization or over-representation of non-informative genomic regions, as well as speeding up computation. For example, one may wish to discard markers when they are in complete (or nearly complete) LD with another, thus bringing the very same information, or selecting markers which are evenly spaced, either physically or genetically, along the genome map.

Molecular data often contain missing value, particularly with the so-called genotyping-by-sequencing (GBS), likely because the fraction of the genome which is re-sequenced is not exactly the same from one individual to another [15]. Most prediction models do not accept missing data, therefore an imputation step is necessary to replace missing values, and various methods have been proposed to achieve the best guess (e.g. fastphase [16]).

Finally, imputed data of (possibly selected) markers are used to predict GEBV using phenotypic observation in a training population. Several methods have been proposed to achieve GEBV prediction. They can be classified into parametric vs semi-parametric methods [17]. In the R environment [18], which is often used in research, several libraries have been specifically developed for genomic selection, such as BGLR [19] or rr-BLUP [20]. However, few of these packages proposes functions to successively achieve the three described steps of marker selection, genotype imputation and model prediction.

The objectives of this manuscript are 1) to describe an integrated software (pipeline) which has been developed from open-source R functions available in various R libraries to enable the three steps to be performed easily and 2) to present an application of this software to carry out genomic predictions using historical data from a bread wheat breeding programme.

## Materials and Methods

### The BWGS pipeline

In the framework of the French flagship programme BreedWheat (www.breedwheat.fr), we developed an integrated pipeline based on R [18] called BWGS (BreedWheat Genomic Selection pipeline). BWGS comprises three modules: 1) missing data imputation, 2) dimension reduction, for reducing the number of markers and/or training individuals and 3) Genomic Estimation of Breeding Values (GEBV) with a choice among 15 parametric and non-parametric methods.

The pipeline comprises two “main” functions (Fig 1)

**Fig 1:**
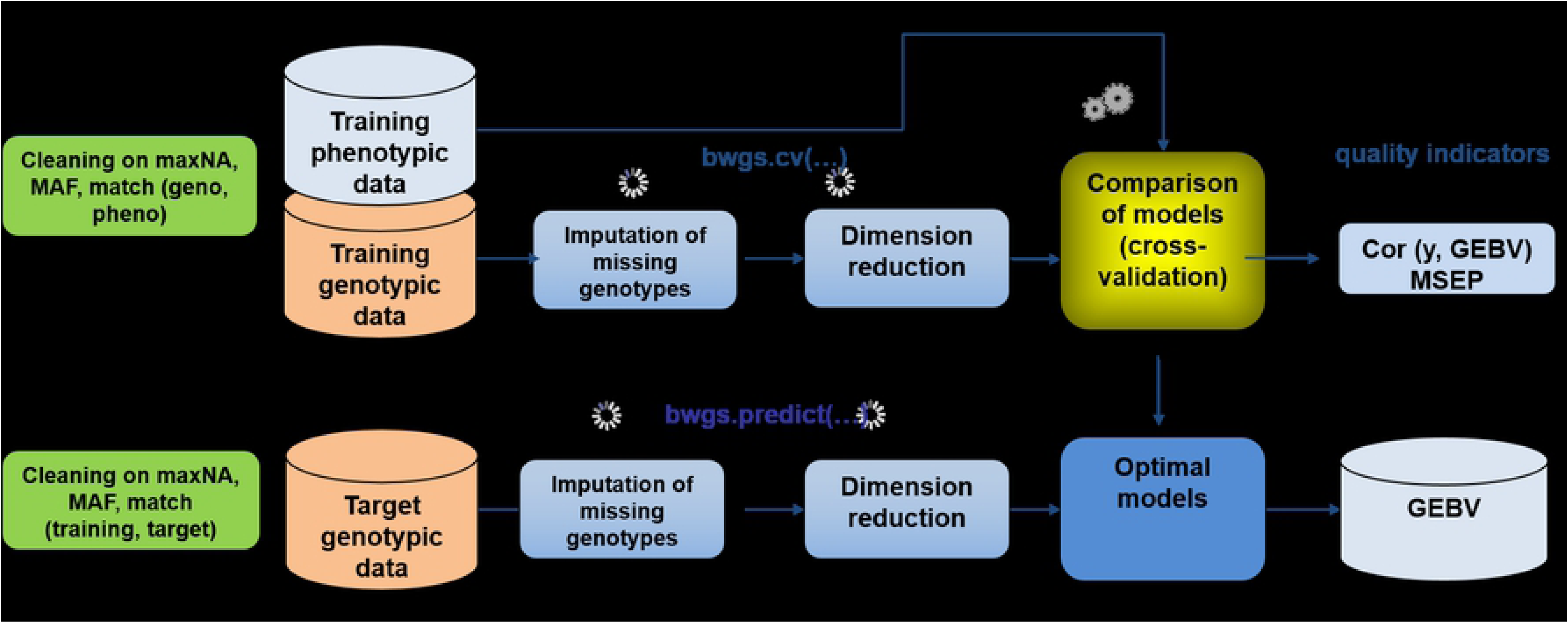
Workflow of the two main functions of BWGS. bwgs.cv does model cross-validation on a training set and bwgs.predict does model calibration on a training set and GEBV prediction of a target set of genotypes. MAF = Minor Allele Frequency, maxNA = maximum % of marker missing data

The first function called bwgs.cv is using both genotyping and phenotyping data from a “training” set or reference population to carry on model calibration and cross validation. Data are randomly split into n “folds”, and n-1 folds are used for training models and predicting the n^th^ one. Computation can be replicated p times, and correlation between GEBV and observed trait are computed for each fold and each replicate, enabling estimates of average and standard deviation of predictive ability (see [21]).

Once the “best” model has been chosen based on quality assessment, a second function, named bwgs.predict is used to build the BEST model using the whole training set (genotyping + phenotyping), then apply the model to the genotyping data of the target population to get GEBV of these new genotypes.

Candidate lines of the target population can then be ranked according to GEBV for single trait (truncation) selection.

Going into more details of the pipeline, the workflow comprises three main steps:

1. A step of (missing) genotyping data imputation. This option can be useful for sources of genotyping data such as GBS. The following options are available:

- MNI: missing data are replaced by the mean allele frequency of the given marker. This imputation method is only suited when there are a few missing values, typically in marker data from SNP chips or KasPAR.
- EMI: missing data are replaced using an expectation-maximization methods described in function A.mat of R-package rrBLUP [20]. This algorithm was specially designed by Poland et al (2012) for the use of GBS markers, which usually give many missing data which are roughly evenly distributed. However, it does not use physical map position, as do other more sophisticated software (e.g. Beagles, [22]). For imputing low density genotyping of a large population to high density available for only a subpopulation, i.e. many markers with many missing data, such software should be used before BWGS.
2. A step of dimension reduction, i.e. reducing the number of markers. This reduction could be necessary to speed up computation on large datasets, depending on computer resources available. The following methods are available

- RMR: Random sampling (without replacement) of a subset of markers. To be used with the parameter “reduct.marker.size”.
- LD (with r2 and MAP): enables “pruning” of markers which are in LD > r2. Only the marker with the least missing values is kept for each pair in LD>r2. To allow faster computation, r2 is estimated chromosome by chromosome, so a MAP file is required with information of marker assignation to chromosomes.
- ANO (with pval): one-way ANOVA are carried out with R function lm on trait “pheno” Every markers are tested one at a time, and only markers with pvalue<pval are kept for GEBV prediction
- ANO+LD (with pval and r2, MAP is facultative): combines a first step of marker selection with ANO, then a second step of pruning using LD option. For research or teaching purposes, an option for randomly sampling individuals has been added, although it is little useful in practical breeding applications. Options for selecting a subset of the training population are:

- RANDOM: a subset of sample.pop.size is randomly selected for training the model, and the unselected part of the population is used for validation. The process is repeated nFolds * nTimes to have the same number of replicates than with cross-validation.
- OPTI: the optimization algorithm based on CDmean [23] to select a subset which maximizes average CD (coefficient of determination) in the validation set. Since the process is long and has some stochastic components, it is repeated only nTimes.
3. A step of model building and cross validation. In the general case of genomic selection, the number of explanatory variables, *i.e.* markers, (largely) exceeds the number of observations, making the classical linear model equation unsolvable. In a review, classified most of the methods that have been proposed to overcome this “big data” problem, into penalized regression (to make them solvable) or semi-parametric methods. Moreover, regression can be solved either analytically as in ridge regression (equivalent to G-BLUP) or iteratively though Bayesian computations. Bayesian methods can differ by the prior density distribution of marker effects, which can be modified boundlessly. In their review [24] describe the main features (e.g. prior…) for 13 methods. The options available for genomic breeding value prediction are:

- GBLUP: performs G-BLUP using a marker-based relationship matrix, implemented through rrBLUP R-library. Equivalent to ridge regression (RR-BLUP) of marker effects.
- EGBLUP: performs EG-BLUP, i.e. BLUP using a “squared” relationship matrix to model epistatic 2×2 interactions, as described by [25] using the BGLR library
- RR: ridge regression, using package glmnet [26]. In theory, strictly equivalent to GBLUP.
- LASSO: Least Absolute Shrinkage and Selection Operator is another penalized regression methods which yield more shrinked estimates than RR. Run by glmnet library.
- EN: Elastic Net [27] which is a weighted combination of RR and LASSO, using glmnet library Several Bayesian methods, using the BGLR library

- BRR: Bayesian ridge regression: same as rr-blup, but Bayesian resolution. Induces homogeneous shrinkage of all markers effects towards zero with Gaussian distribution [24].
- BL: Bayesian LASSO: uses an exponential prior on marker variances priors, leading to double exponential distribution of marker effects [28].
- BA: Bayes A uses a scaled-t prior distribution of marker effects [6].
- BB: Bayes B, uses a mixture of distribution with a point mass at zero and with a slab of non-zero marker effects with a scaled-t distribution [29].
- BC: Bayes C same as Bayes B with Gaussian a distribution for non-zero marker effects[19] A more detailed description of these methods can be found in (http://genomics.cimmyt.org/BGLR-extdoc.pdf.) Four semi-parametric methods

- RKHS: reproductive kernel Hilbert space and multiple kernel MRKHS, using BGLR [30–31]. Based on genetic distance and a kernel function to regulate the distribution of marker effects. This methods is claimed to be effective for detecting non additive effects.
- RF: Random forest regression, using randomForest library [32]. This method uses regression models on tree nodes which are rooted in bootstrapping data. Supposed to be able to capture interactions between markers.
- SVM: support vector machine, run by e1071 library. For details, see LIBSVM: a library for Support Vector Machines https://www.csie.ntu.edu.tw/~cjlin/papers/libsvm.pdf
- BRNN: Bayesian Regularization for feed-forward Neural Network, with the R-package BRNN [33]. To keep computing time in reasonable limits, the parameters for the brnn function are neurons=2 and epochs = 20.

The criteria used to estimate model’s quality are 1) the Pearson correlation between adjusted phenotype and GEBV, computed over the whole data set (i.e. by merging the value from the nFolds) and 2) the square-root of the mean-squared error of prediction RMSEP. These criteria are provided for each replicate (nTimes) as well as mean and standard deviation over replicates. A table summarizes phenotype, estimated breeding value and its standard deviation, as well as the coefficient of determination for each individual GEBV computed as CD=sqrt(1-(SD(GEBV)^2^/VAR(GEBV)), when the package used does provide estimate of variance and standard deviation of GEBV, as it is the case for BGLR.

### Application to real wheat breeding: plant materials and phenotypic data

A training population was built by gathering historical data from the INRA-Agri-Obtentions breeding programmes. To obtain robust phenotypes, we kept data from lines which have been evaluated in multisite trials (> 6 locations), usually for 1-2 years. They were developed from inbred lines crosses, followed by 6-8 generations of self-pollination. Few selection is made during the first 3 generations (i.e. F2 to F4), which are harvested in bulk, then visual selection on simple traits is made on plants or rows at generations F4 and F5. Selected F5 families are multiplied in field plots, which allow a rough evaluation of yield. Then enough seeds are available and selected lines are put into the evaluation network, usually at the F7 generation, when breeding lines are nearly homozygous. Typically, about 300-400 pair crosses are made each year, 120-150 F7 lines are put into first year multisite trials, then 50-60 are evaluated for a second year, and the very happy few (1-3) are then entering the official registration trials in France, to hopefully become a commercial variety, 10-12 years after the initial crosses. On the whole, our database for field data had 77,176 records on 1715 genotypes, *i.e.* 45 single plot measurements per genotype, spread over 15 years (2000-2014), 10 locations in France and two managements (high *vs* low inputs). Of course these data are highly unbalanced, each genotype being evaluated, on average, during 1.5 years in 7.60 environments (site x year) with usually two replicates by management, and most connections in the design are between two successive years, with a few control varieties (usually four in each trial) being evaluated on a longer period. These figures are for grain yield, the most documented trait. The whole dataset is described in [34] and can be recovered at urgi.versailles.inra.fr (INRA Small Grain Cereals Network Phenotypic Trials dataset). To illustrate the use of BWGS, we will concentrate on grain yield in high input management, i.e. an estimate of the genetic and climatic potential in each environment.

These highly unbalanced raw data require a pre-treatment step to correct for non-genetic factors as much as possible. The following mixed model was applied to estimate “corrected” genotypic means (BLUE) using the mixed model in R (library lme4):

Y=lmer (yield∼geno + (1|year:site:trial:block) + (1|year:site:geno)), where : stands for nested effects, e.g. block within trial within year.

Where geno being the main genotypic effect (fixed) and all other effects being considered as random effects to be corrected for. Note that these genotypic “BLUE” are highly correlated (r=0.94) with BLUP estimates (*i.e.* when genotypes are also considered as random effects with identity matrix for modelling covariances). But BLUE are not shrinked, and thus can be more easily used in a second step involving mixed modelling.

The same model with geno as random effect was used to estimate variance components and rough estimate of broad sense heritability as σ ²_g_ / (σ ²_g_ + σ ²_ge_ /nenv + σ²_e_ /nplot) σ

Where σ²_g_ is the genotypic variance component, σ²_ge_ is the GxE variance component (i.e. year:site:geno) and σ²_g_ is the residual variance. nenv is the average number of environments per genotype and nplot the average number of plots per genotype in the dataset used.

Out of these “HQ-phenotyped” lines, 760 were also genotyped with the TaBW280K SNP chip [35] and used for testing BWGS. After quality check, a matrix of 188,406 high quality, polymorphic markers was finally used. To illustrate BWGS pipeline, we used a subset of 60,912 markers with consolidated map position.

Cross Validation used randomly sampled 90% of the genotypes as the training set and the remaining 10% genotypes as the “validation set”. The resampling process was iterated 100 times to estimate an empirical mean and standard deviation. Since the true breeding value was unknown, predictive ability (PA) was measured by the Pearson correlation between GEBVs and the adjusted phenotypic values (BLUE) across all folds, i.e. the Hold accuracy, which is supposed to be less biased that a fold-by-fold estimate [36].

## Results

### Summary statistics of the breeding population

Genotypic data have been used to estimate the additive relationships matrix according to [37] with the A.mat function of the R package rrblup. After scaling on a 0-1 scale (1 on diagonal), the values can be regarded as estimates ofcoancestry coefficients, whose distribution is shown in Fig 2a. Omitting the diagonal, coanscestry coefficients range between 0.05 and 0.5, with a median around 0.18, i.e. slightly less than the value expected for half-sibs. The heat map of Fig 2b displays no clear-cut structure into distinct groups, although some groups along the diagonal are made of lines that are more related than average. Such absence of structure allows the use of random sampling cross-validation. A plot of principal component analysis of Euclidian distance matrix among the 760 lines (Fig 2c, R command cmdscale) again does not show any clear structure according to the year of first evaluation, thereby justifying again the use of random cross-validation over years.

**Fig 2:**
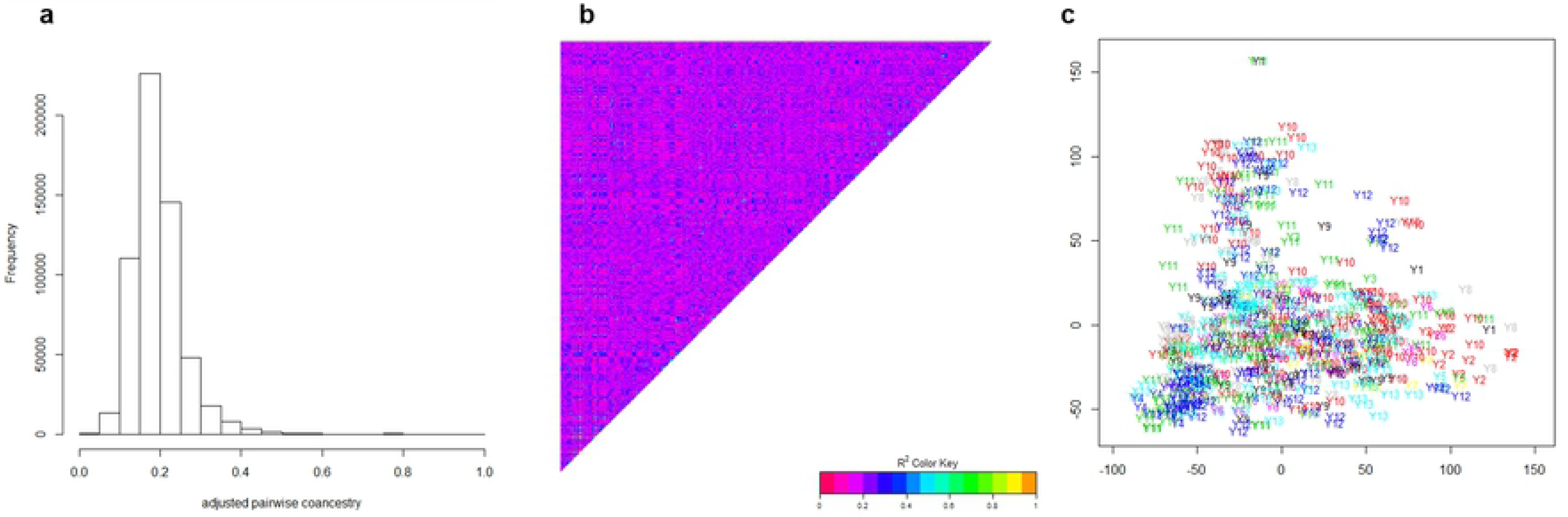
Histogram (a), heat map (b) and PCA plot (c)of the scaled coefficient of coancestry between the 760 breeding<u>. Yx represents the year of first evaluation of a given line.</u>

### Imputation of missing data

The first step of data cleaning with MAF=5% and maximum marker missing data = 20% led to retain 47,839 markers out of 60,912. Since adjusted Yield BLUE were complete for the 760 genotypes, the final dimension of geno matrix was 760 x 47,839. This SNP dataset contained on average 0.93% of missing data. Given the small proportion of missing data, no difference in predictive ability was observed between the two imputation methods. To test the efficiency of imputation methods on prediction accuracy, we generated new genotypic matrices with 20%, 40%; 60% and 80% of missing data, randomly distributed in the dataset. Imputation by mean allele frequency took 0.15 minute, while imputation by the EM algorithm of A.mat took around 15 minutes, whatever the proportion of missing data. Predictive ability for grain yield from imputed dataset using the two methods is presented in Fig 3. The two methods give similar results for up to 20% missing Data, then imputation by EMI allows higher predictive abilities than imputation by the mean allele frequency. When 80% missing data are randomly generated, the predictive ability of EMI-imputed set is still 0.418, while it drops to 0.329 with the MNI-imputed set.

**Fig 3:**
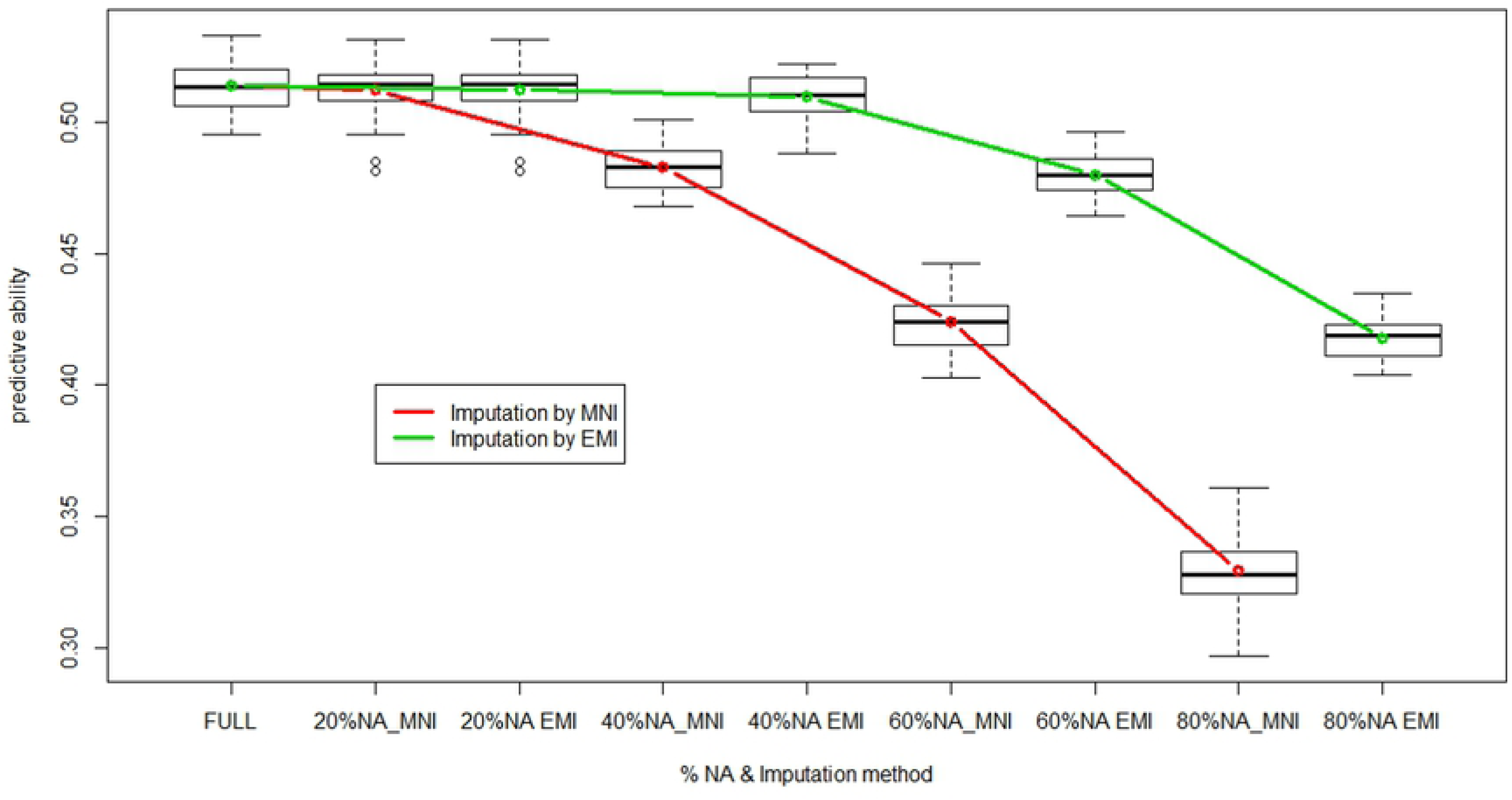
Predictive ability of GBLUP as a function of % of randomly generated missing data, with two imputation methods 1) mean allele frequency or 2) expectation-maximization (EM in A.mat function of rrBLUP library)

### Effects of sampling markers

Table 1 shows the number of markers selected by either one-way ANOVA at different pvalue thresholds, or by pruning markers with LD > threshold values from 0.5 to 0.98. For ANOVA selection, two strategies have been used, namely 1) GWAS was carried out only once using the whole dataset, i.e. training + validation lines and 2) GWAS was carried out using training lines only, i.e within each time x fold replicate. Predictive abilities achieved with the selected marker subsets are also shown in Table 1.

**Table 1:** Number of markers and predictive abilities achieved with marker subset from 1) random selection (RMR option); 2) GWAS on whole marker set (option ANO); 3) GWAS on training set only (not included) and 4) LD-pruning.

| random selection |  | GWAS |  | GWAS whole set | GWAS training set | LD pruning |  |  |
| --- | --- | --- | --- | --- | --- | --- | --- | --- |
| Nb markers | Pred ability | Pvalue | Nb markers | Pred ability | Pred ability | LD_threshold | Nb markers | Pred ability |
| 47739 | 0.525 | 1.0 | 47839 | 0.526 | 0.526 |  |  |  |
| 25000 | 0.52 | 0.05 | 14390 | 0.573 | 0.528 | 0.98 | 15773 | 0.538 |
| 10000 | 0.515 | 0.01 | 8325 | 0.579 | 0.529 | 0.95 | 9297 | 0.541 |
| 5000 | 0.514 | 0.001 | 4000 | 0.596 | 0.519 | 0.9 | 6661 | 0.54 |
| 2000 | 0.505 | 0.0001 | 1820 | 0.614 | 0.515 | 0.8 | 4462 | 0.544 |
| 1000 | 0.493 | 0.00001 | 806 | 0.576 | 0.473 | 0.7 | 3234 | 0.536 |
| 500 | 0.447 | 0.000001 | 323 | 0.551 | 0.438 | 0.6 | 2376 | 0.527 |
| 200 | 0.392 |  |  |  |  | 0.5 | 1704 | 0.521 |
| 100 | 0.299 |  |  |  |  |  |  |  |

Results are also displayed in Fig 4.

**Fig 4:**
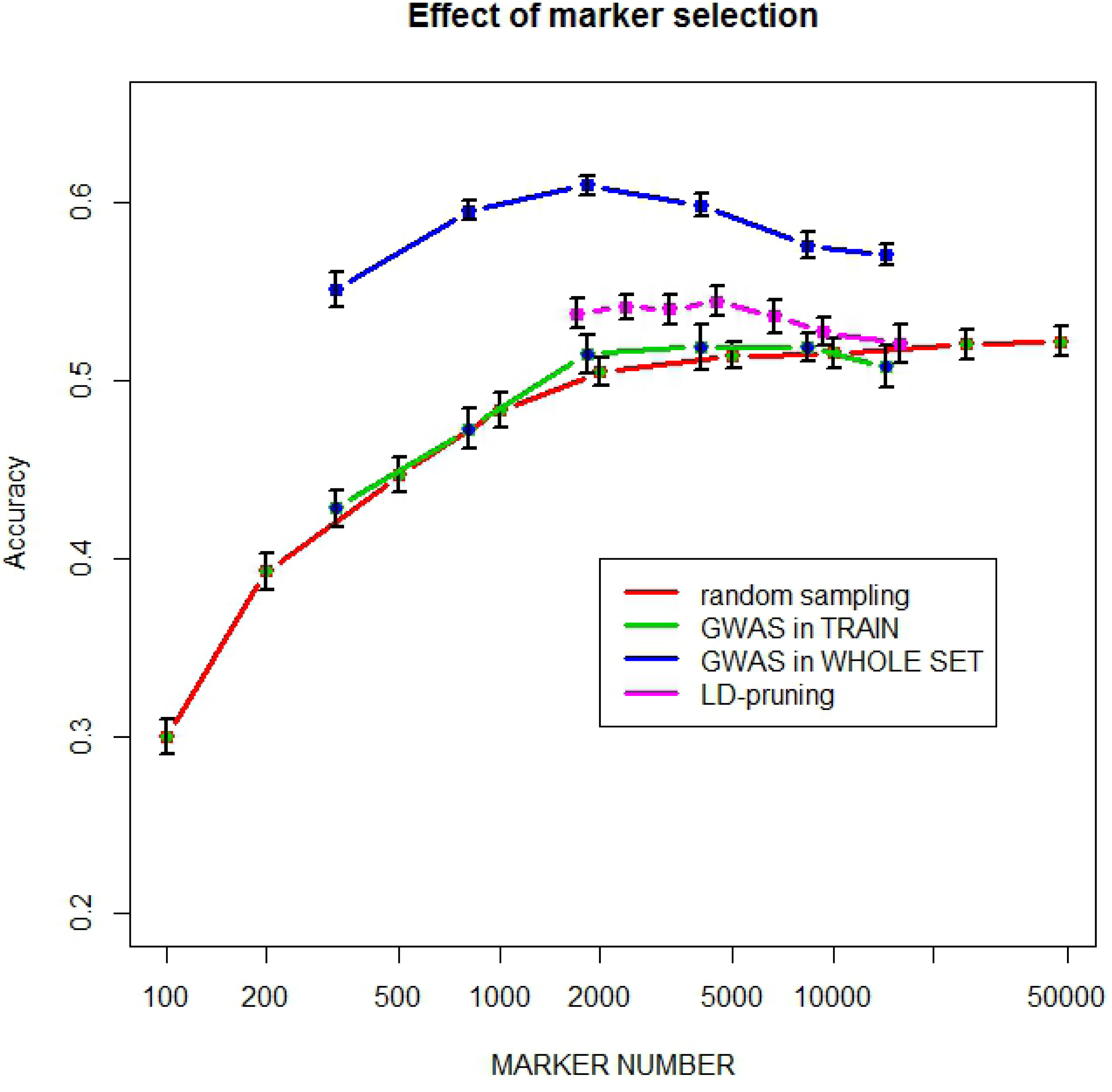
Predictive ability as a function of marker number selected either 1) randomly, 2) by GWAS in the training set within each replicate* fold, 3) by GWAS on the entire population and 4) by LD-pruning.

As expected, predictive ability increases with the number of randomly sampled markers, with a steady plateau above around 5000 markers and a best predictive ability of 0.525. This clearly shows that the extent of LD is large enough in such elite breeding lines to allow most QTLs for yield being captured with 5000-10000 markers, i.e. on average one every 1.7 Mbase. This is illustrated by marker selection based on LD-pruning, i.e. removing markers which are in LD > threshold with any other marker. The marker with the least number of missing value is conserved, otherwise the choice is random. The highest value of 0.544 is obtained with 4462 markers with pairwise LD >0.8. When marker selection is based on ANOVA carried out on the entire population, there is an optimum predictive ability of 0.614 achieved with 1820 markers (pvalue = 0.0001). However, these results were only found when marker selection is made only once by association analysis on the whole dataset, which is then split into a training and a validation set. When marker selection is made within the loop only in the training set, then the predictive ability of the validation set is only slightly improved for N= 2000 and 5000 markers, compared to random sampling, but never exceeds the predictive ability achieved with the whole set of markers. This clearly illustrates that overfitting does occur in the first case.

### Effect of different calibration set sampling strategies on prediction accuracy

For this study, we used the whole set of 47,839 markers; which gave a predictive ability of 0.525 in 10-fold cross validation using the complete dataset of 760 breeding lines, i.e. with a training set of 684 lines and a validation set of 76 lines. Predictive ability was estimated with GBLUP using calibration set of size CS= 50, 100, 200, 300, 400, 500, 600 and 700, validation being made using the remaining lines. Fig 5 shows the increase of both predictive ability and computing time as the size of training set increases. Computing time is given as the proportion of that needed for 100 replicates x 10 folds with TS = 700, which was roughly 60 minutes. As expected, predictive ability does increase with the size of calibration set, while its variability decreases, with the notable exception of TS = 700. This may be due to the small size of the validation set, which is only 76 with TS = 700, making correlations between phenotype and GEBV more erratic than those observed with larger validation sets.

**Fig 5:**
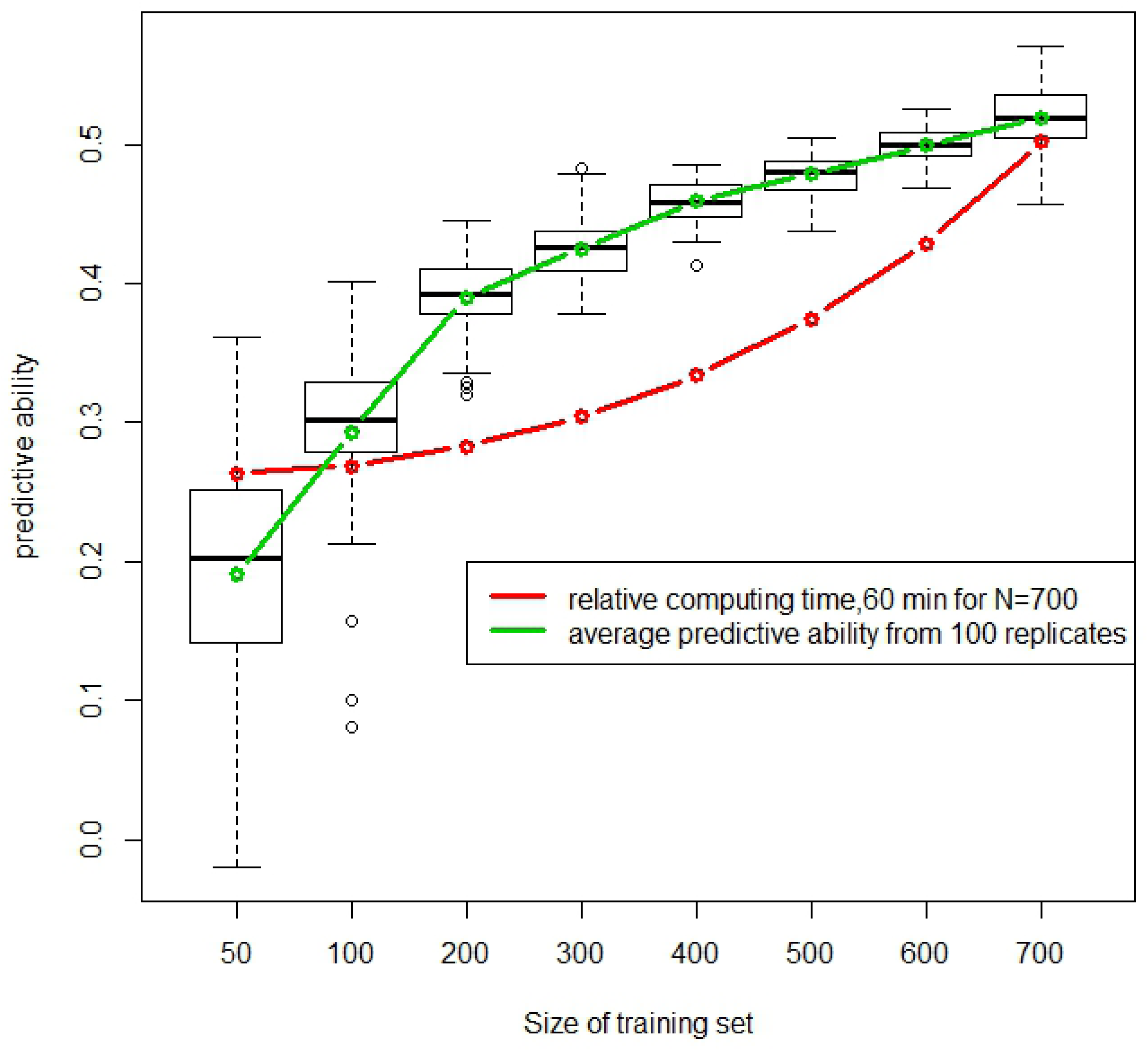
Distribution of predictive ability according to the size of randomly selected training population and relative computing time.

Two strategies were compared as illustrated in Fig 6: 1) the random sampling among the 760 lines as in Fig 5 and 2) selecting an optimized subset with the CD-mean criteria as described by [23]. Although this algorithm should be deterministic and give always the same subset, some stochasticity remains in the drop-replacement procedures, which explains that predictive ability still have some residual variation as illustrated by the error-bar. Fig 6 shows that the optimization algorithm does improve predictive ability for small-medium size of the training set compared to random sampling. as predicted by the theory and already reported [23]. This advantage disappears when the proportion of sampled individuals increases.

**Fig 6:**
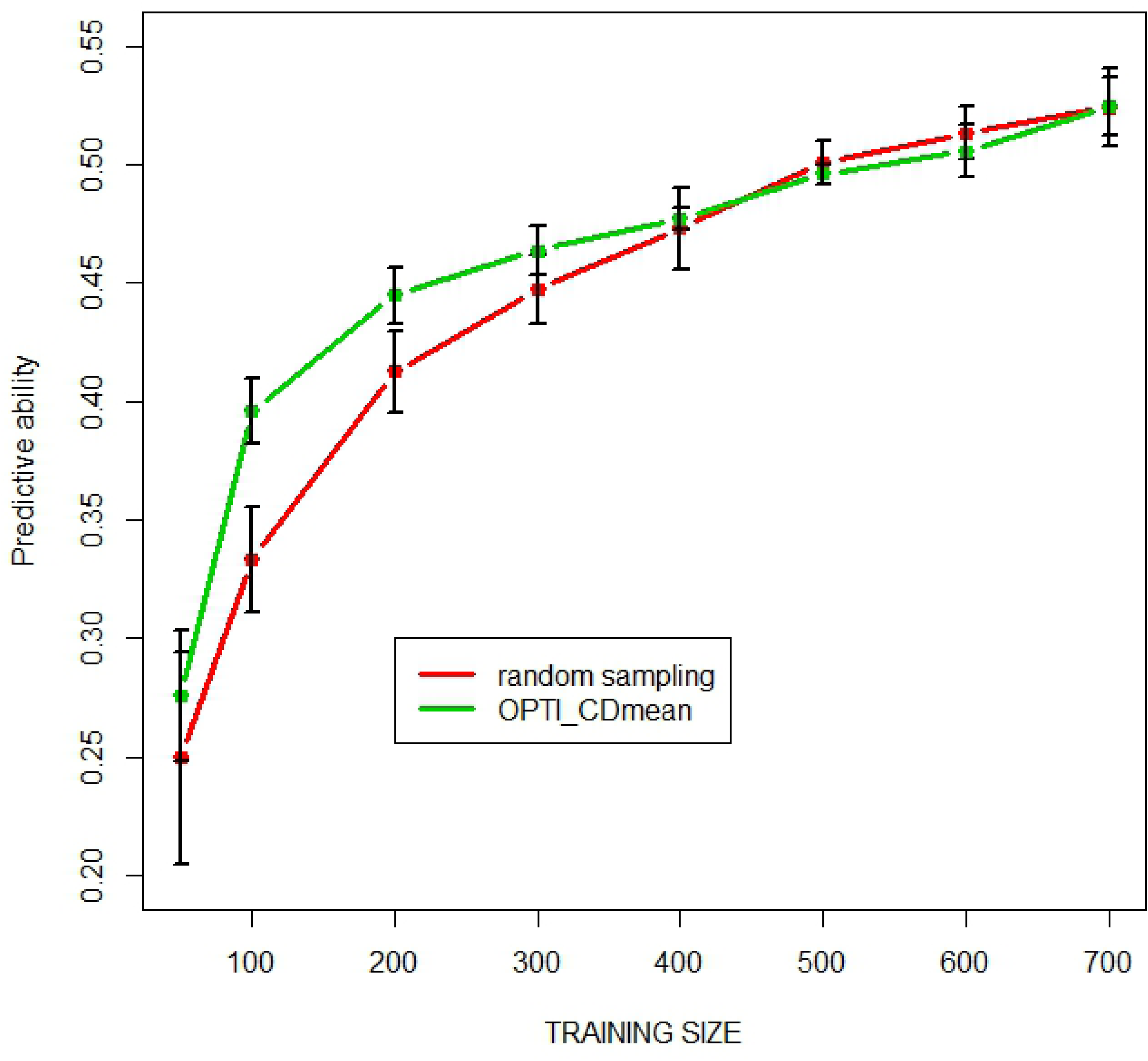
Predictive ability as a function of training population size selected either 1° randomly or 2) by the CD-mean optimization algorithm of [23]

### Efficiency of various prediction methods

With again the whole set of 47,839 markers, we used all prediction methods available in BWGS to estimate prediction accuracy in 100 independent 10-fold cross validations. Computing time varied considerably from one method to another, as illustrated in Fig7. BRNN is, by far, the most demanding methods, with 2214 minutes (nearly 4 days) needed to carry out 100 replicates of 10-folds cross validation, while the least demanding is GBLUP with 28 minutes. Of course these values must be taken only for comparison, as they are highly dependent on the computer characteristics. Note that EGBLUP, which is an extension of GBLUP with epistatic interactions being modelled by the product of additive relationship matrix with itself, takes 344 minutes instead of 28. The other computationally intensive methods are multi-kernel RKHS (637 minutes) and random Forest RF (382 minutes).

In Fig 7, we discarded results from SVM methods, which gave very poor predictive ability (0.25), although in a reasonable time of 31 minutes. Predictive abilities of the other 14 methods range from 0.475 to 0.543, with no relationship with computing time, which can be considered as an estimate of method complexity. Note that the support vector machine SVM compared with other methods when 5000 random markers are used, but seemed to be unable to deal with 47,000 markers. LASSO and elastic net (from glmnet library) and BRNN give the worse predictive abilities around 0.48, while random forest regression RF gives the highest of 0.543, slightly above that of the reference GBLUP (0.525). Note that the three methods that outperform GBLUP are thought to take into account non additive marker effects. However, in this practical case of grain yield prediction, they did not show a dramatic advantage over GBLUP. BRNN is another model-free, machine learning method, often supposed to give more accurate prediction than linear regression methods. The relatively poor predictive ability observed in this study can be caused by insufficient computer resources allocated. Other parameterization (e.g. number of neurone layers, epochs…) may have given better predictive abilities.

**Fig 7:**
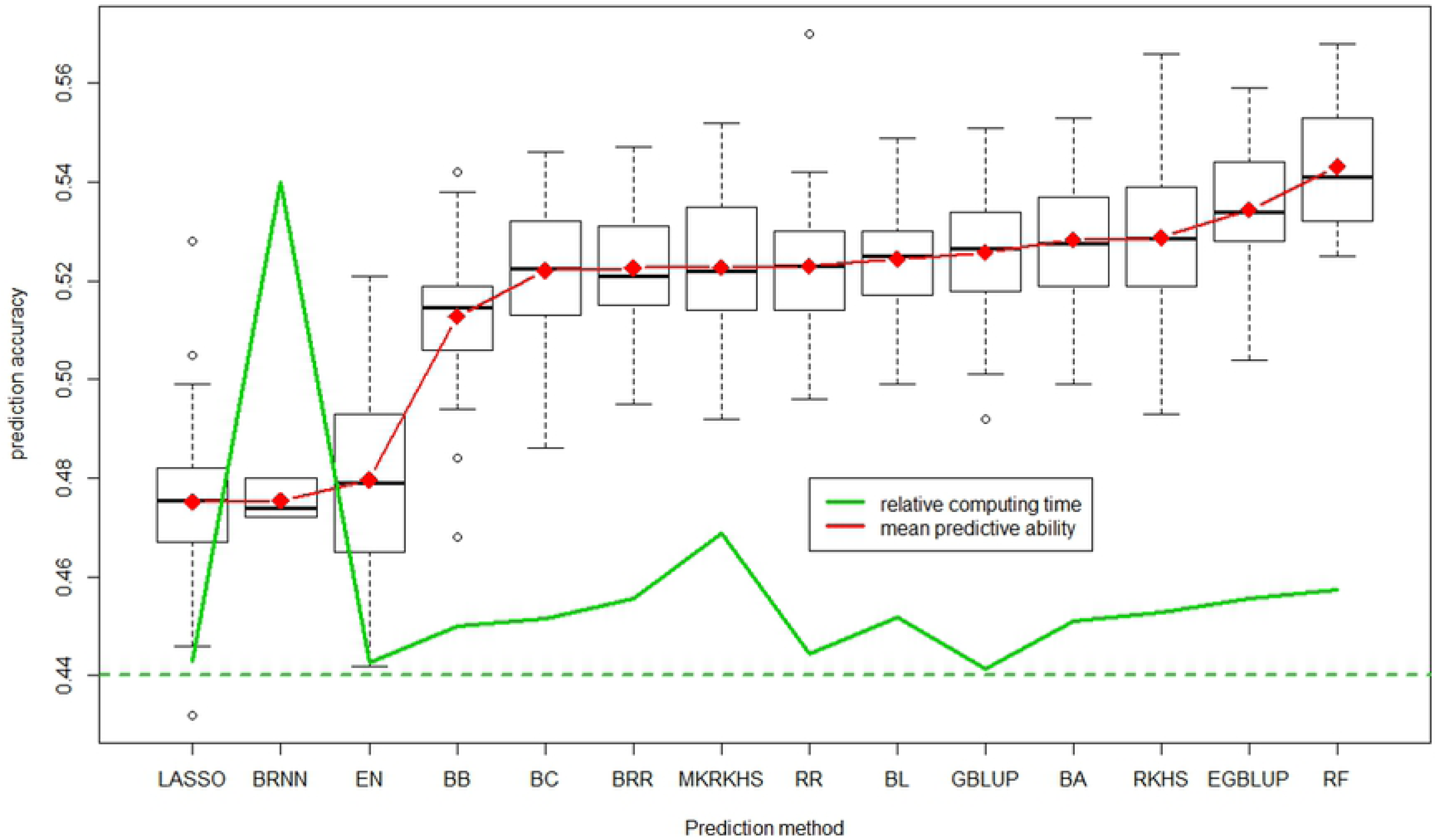
Distribution of predictive ability of the 100 replicates for each of the 14 methods which worked correctly, average is in red and relative computing time in green line.

Perhaps as important as predictive ability is the consistency of GEBV estimated by different methods. Fig 8 shows the correlation between GEBV (averaged over the 100 replicates) predicted by the 14 successful methods (omitting SVM). The minimum correlation of 0.85 is between random Forest and Bayesian ridge regression, while many pairwise correlation are close to 1. When omitting RF, which is the method whose prediction are least related to the others, all correlations are above 0.92, thus all methods can be considered as giving highly consistent prediction of GEBV.

**Fig 8:**
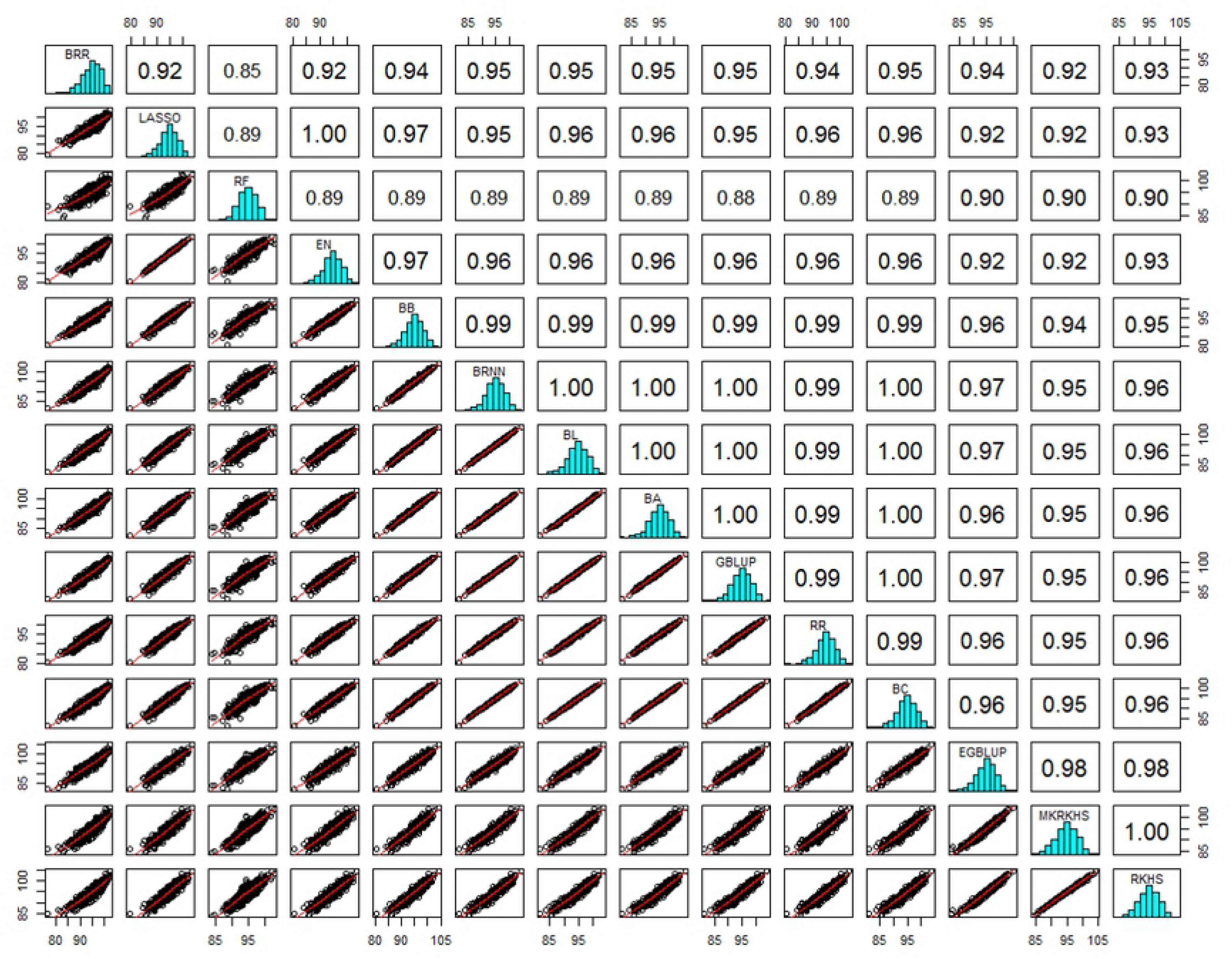
Histograms, bi-plot and correlation values among predicted GEBV obtained with 14 methods.

## Discussion

Genomic selection programmes are now routinely used in dairy cow breeding, and benefit from huge phenotypic data recorded in past years on milk production of thousands of females, usually related by well-known pedigree relationships (i.e. mother and father known without error). This is however not yet the case in other species like sheep, although effort are being developed in so-called minor species. Therefore, using a relatively cheap genotyping, dairy cow breeders usually have very large population for training GS models, which led to highly accurate predictions. In many animal studies, the oldest animals are used as training set and the youngest as validation set.

Contrasting with animal breeding, most plant breeding programmes do not have very large population sizes, although each breeding company manipulates hundreds or thousands of candidates, since companies are usually reluctant to share and merge datasets. Breeding companies have developed in-house biostatistical tools for calculating GEBV with semi-automated pipeline, since time is often short between data production (e.g. grain harvest) and selection decision (e.g. sowing next generation). There are also several publically available tools which have been developed, particularly as R libraries. Among the most popular we can mention glmnet [26], BGLR [19], rrBLUP [20] or Synbreed [38]. Recently, integrative packages have been developed, which rely on existing public R-libraries. An example of such packages are G2P and the one presented here called BWGS. G2P [39] proposes 17 prediction methods from 10 R-libraries, among which BGLR, glmnet and randomForest as in BWGS. As BWGS, there are also two main functions, G2PCrossValidation and G2P to apply prediction model to a test set with only marker data. Compared to BWGS, G2P offers much more options for tuning parameters of the numerous functions/libraries called by the two main functions. As a drawback, handling G2P appears more complex, and its complete use requires reading the notices of original libraries, since the notice of G2P does not provide enough details on the possible parameters and options. Moreover, although G2P does content a quality control module, it is not directly integrated into the main function as in BWGS. Comparatively, in BWGS, we have chosen to fix most internal parameters with defaults values, which have been tested to be adapted for medium size datasets as provided in the example (47 K SNP, 760 training lines), while maintaining computing time into reasonable limits for desktop computers. It is therefore easy to use, especially for beginners, and of course parameters can be modified quite easily in the source code to adapt larger datasets.

In our highly unbalanced breeder’s dataset, when discarding control lines which were regularly replicated over years, the average number of environment per studied lines is 10.4, with two replicates by environment (site x year). This led to an estimate of broad sense heritability of 0.76 and therefore a theoretical upward limit of prediction accuracy of 0.872. When using random cross validation and the whole set of 47,839 markers, the achieved predictive ability is 0.525, which would correspond to an accuracy of 0.603 according to formula in [21]. However we do not fully trust in this formula, since heritabilities are often poorly estimated. We do prefer keeping predictive ability as a criteria for comparing models and strategies.

Our results compare with previously reported predictive ability estimated through random cross-validation, for example 0.36-0.53 [40] and 0.32-0.59 [41] for grain yield in bread wheat with training population of a few hundreds lines, and up to 0.65 with a training set of 2325 elite European wheat lines [42]. The use of random cross validation seems to be justified, since no clear-cut structure appears in the set of lines. In particular, lines put into trial in a given year are not more related with each other than with lines put into trails another year. Then the predictive ability obtained in random cross-validation should be valid for any other set of lines showing a similar degree of relatedness than within the training set.

### Effect of marker density and training population size

Although GS theory has been elaborated to cope with the over parameterization problem (number of markers >> number of observations), it empirically appears that adding more markers than needed does not improve predictive ability. This was already observed in many reports. Among other, a figure similar to our Figure 3 can be found in [43]. In this empirical study in maize, GS accuracy reach a plateau with 7000 randomly selected markers in a “natural” population, and with only 2000 markers in biparental populations. In a recent simulation study with high density coverage, the same authors even stated that the accuracy obtained using all SNPs can be easily achieved using only 0.5 to 1.0% of all markers [44]. This clearly illustrates that, once every QTL information is captured by one marker in LD, adding more markers is useless. This of course relates to the average linkage disequilibrium between adjacent markers. In our study, the material is made of hundreds of related families, each of small size, and the plateau is reached around 2000-5000 random markers, a value close to that observed for maize natural population, while [45] reported that 256 markers were enough to achieve maximum accuracy in wheat bi-parental populations. Similarly, in a population of 235 soybean varieties, predictive ability did not change much, whatever the number of markers (ranging 200-5200) and the way they were selected, either at random or one per haplotype block [46]. In a study of wheat breeding lines [47] found a plateau for predictive ability of yield around 2000 random markers as in the present study. Avoiding selection of markers pairs which are in high LD (LD-pruning) further improves predictive ability compare to random sampling. This was reported in a soybean study, in which the authors found a 4% increase of prediction accuracy when selecting markers from haplotype blocks rather than random or equidistant.

Selecting markers that are significantly associated with QTLs can achieve higher predictive ability than randomly selected markers, and surprisingly even higher predictive ability than using all markers. However, it is clear from Fig 4 that selecting markers from their Pvalue in GWAS carried out on the whole population; i.e. including validation lines, does led to overfitting and this approach must be avoided and cannot be used in practice. This was reported many times. For example, in a wheat study [47] selected markers by GWAS on the training set only, as we did also in Fig 4. However, they observed a gain in predictive ability of up to 0.2, particularly for very small number of markers (<100), while we only had small improvement of about 0.01 with a maximum gain for 2000 markers. Nearly similar results were reported by [48].

In theory, the prediction accuracy is positively related to the training population size, as established by simulation studies [49–51]. This was confirmed in many empirical studies such as those already mentioned [42, 46, 47]. As expected from the theory, optimizing a subset of training lines gave higher PA than random selection. This was confirmed in empirical studies [23] then [52–53] in the case of population structure. Their optimization algorithm uses a simulated annealing approach. Other optimization methods have been proposed, such as a genetic algorithm [54] and also led to higher PA for a given training size.

### Effect of prediction method

Prediction methods have been classified into parametric and non-parametric (or semi-parametic) methods, and parametric methods sometimes split into penalized approach and Bayesian approach (for reviews see [17] and [55–57]). In our study, most methods gave close values of predictive abilities, ranging 0.475 −0.543, with the noticeable exception of SVM, which worked with up to 5000 markers (data not shown), but failed with the 47 K markers. Some methods seem to give poorer results, such as LASSO and EN (elastic net) from the glmnet library, and also BRNN. In this latter case it may be due to insufficient computer resources, who led us to use restrictive parameters (e.g. number of iterations, number of neurone layers…), thereby limiting performances of this highly demanding method. In any case predictive ability is not related to computing time, with the less demanding GBLUP ranking the fifth best method.

Although there is no clear-cut separation between parametric vs non-parametric methods, nor between penalized vs Bayesian approach, it seems that methods that are supposed to better capture non additive and/or non-linear effects such as EGBLUP or RKHS gave higher PA, as already reported [33,42,48,56]. Low difference in predictive ability has already been reported by [39] who analysed three wheat populations for GEBV using 10 statistical models, with RKHS being the most accurate and Support Vector Machine the least accurate methods, as we also found in our study.

In a simulation study, [58] showed that with 121 markers with additive effects, RKHS and radial basis neural network (close to our BRNN option) clearly out passed the linear Bayesian LASSO, but it was no longer the case when adding the 7260 interactions between markers. The authors stated that “adding non-signal predictors can adversely affect the predictive accuracy of the non-linear regression models”. Other studies have shown the interest of machine learning methods for genomic prediction [59–60]. In our study, this was not the case for RKHS, but for SVM and BRNN. It may have been better to choose the set of LD-pruning selected markers to optimize predictive ability of these methods.

But perhaps more important than the predictive ability is to know whether the different methods give similar values or at least similar rankings of GEBV for candidates. Indeed correlations between predicted GEBV with the 15 methods range from 0.85 to nearly 1. The highest values being observed between linear parametric methods, whatever based on penalized regression or Bayesian approaches. Machine learning methods such as SVM and BRNN are least related to others, except with MKRKHS. Given the close values of both predictive abilities and GEBV estimates among methods, it seems reasonable to keep the historical GBLUP as a reference method, for its simplicity and fastness, at least for polygenic traits such as grain yield chosen to illustrate this study.

### Conclusion

The R pipeline we have developed is based on publically available libraries and therefore offers a full freedom to operate. It is easy to handle and allows a wide range of options for missing data imputation, marker or training set selection and prediction methods. Its parameterization was fixed for medium sized datasets to make it easy to use for beginners or teaching. Applying this tool with defaults parameters to a set of elite breeding lines with historical data from yield trials allowed us to obtain similar results to those reported on other wheat populations. The options for subsampling markers and/or training set enabled us to illustrate theoretical expectations (e.g. [61]). The predictive abilities obtained on this population of limited size are encouraging for the success of genomic selection in applied wheat breeding. Of course, BWGS does not deal with all challenges. It is now admitted that most methods give reasonably high accuracy, although recent studies claim that prediction accuracy could be improved with new alliances to share data across breeding programmes [62]. Rather, the challenges for future wheat breeding are 1) efficient implementation in real breeding schemes and/or adapting selection schemes with step(s) of GS (e.g.[63–64]), 2) prediction of GxE (e.g. [65]for review) and 3) incorporating multitrait selection (e.g. [66]).

Future developments of BWGS are ongoing to address these challenges, particularly multitrait selection.

The source code of BWGS R functions as well as example files and notice are available on https://forgemia.inra.fr/umr-gdec/bwgs

## Acknowledgements

This work was supported by the BreedWheat project thanks to funding from the French Government managed by the National Research Agency (ANR) in the framework of Investments for the Future (ANR-10-BTBR-03), France AgriMer and the French Fund to support Plant Breeding (FSOV).

## References

1. Lande R, Thompson R (1990) Efficiency of Marker-AssistedSelection in the Improvement of Quantitative Traits. Genetics 124:743–756

2. Hospital F, Moreau L, Lacoudre F, Charcosset A, Gallais A (1997) More on the efficiency of marker-assisted selection. Theoretical and Applied Genetics 95:1181–1189

3. Bernardo R, Moreau L, Charcosset A. Number and fitness of selected individuals in marker-assisted and phenotypic recurrent selection. Crop Sci 2006;46: 1972–1980.

4. Moreau L, Charcosset A, Hospital F, Gallais A (1998) Marker-assisted selection efficiency in populations of finite size. Genetics 148:1353–1365

5. Whittaker, J.C., R., Thompson, M.C. Denham. 2000. Marker-assisted selection using ridge regression. Genet. Res. 75:249–252.

6. Meuwissen THE, Hayes B, Goddard ME (2001) Prediction of total genetic value using genome-wide dense marker maps; Genetics 157:1819–1829

7. Goddard ME, Hayes BJ (2007) Genomic selection. J Anim Breed Genet 124:323–330.

8. Schaeffer, L. R. (2006) Strategy for applying genome-wide selection in dairy cattle. Journal of Animal Breeding and Genetics 123: 218–223 DOI: 10.1111/j.1439-0388.2006.00595.x

9. Bernardo R, Yu JM. Prospects for genomewide selection for quantitative traits in maize. Crop Sci 2007;47:1082–1090.

10. Heffner EL, Sorrells ME, Jannink JL (2009) Genomic Selection for Crop Improvement. Crop Science 49:1–12

11. Crossa J., de los Campos G., Perez P., Gianola D., Burgueño J., Araus J.L., Makumbi D., Singh R.P., Dreisigacker S., Yan J, Arief V., Banziger M. and Braun H.J. 2010. Prediction of Genetic Values of Quantitative Traits in Plant Breeding Using Pedigree and Molecular Markers. Genetics 186: 713–724

12. Jannink J. L.; Lorenz A. J.; Iwata H. (2010) Genomic selection in plant breeding: from theory to practice. Briefings in Functional Genomics & Proteomics 9: 166–177

13. Iwata H.; Jannink J. L. (2011) Accuracy of genomic selection prediction in barley breeding programs: a simulation study based on the real single nucleotide polymorphism data of barley breeding lines. Crop Sci 2011;51: 1915–1927

14. Lorenz AJ, Chao S, Asoro FG, Heffner EL, Hayashi T, Iwata H, Smith KP, Sorrells MK and Jannink J L (2011) Genomic selection in plant breeding: knowledge and prospects Adv Agron 110, 77–123

15. Poland, J., Endelman, J.B., Dawson, J., Rutkoski, J.E., Wu, S., Manès, Y., Dreisigacker, S., Crossa, J., Sanchez-Villeda, H., Sorrells, M.E., & Jannink, J. (2012). Genomic Selection in Wheat Breeding using Genotyping-by-Sequencing. The Plant Genome 2012;5:103–113. doi: 10.3835/plantgenome2012.06.0006

16. Scheet, P and Stephens, M (2006). A fast and flexible statistical model for large-scale population genotype data: applications to inferring missing genotypes and haplotypic phase.Am. J. Hum. Genet. 2006; 78:629–644

17. Desta, Z.A., Ortiz, R. (2014) Genomic selection: genome-wide prediction in plant improvement. Trends Plant Sci. 2014 Sep;19(9):592–601. doi: 10.1016/j.tplants.2014.05.006. Epub 2014 Jun 23.

18. R Development Core Team, 2011 R: A Language and Environment for Statistical Computing. R Foundation for Statistical Computing, Vienna. http://www.R-project.org

19. Pérez P, de los Campos G. Genome-Wide Regression and Prediction with the BGLR Statistical Package. Genetics. 2014;198(2):483–495. doi:10.1534/genetics.114.164442.

20. Endelman J.B. (2011) Ridge Regression and Other Kernels for Genomic Selection with R Package rrBLUP. The Plant Genome 4:250–255

21. Estaghvirou SBO, Ogutu JO, Schulz-Streek T, Knaak C, Ouzunova M, Gordillo A, Piepho AP (2013) Evaluation of approaches for estimating the accuracy of genomic prediction in plant breeding. BMC Genomics 12:860 http://www.biomedcentral.com/1471-2164/14/860

22. Browning BL, Browning SR. Genotype Imputation with Millions of Reference Samples. American Journal of Human Genetics 2016;98(1), 116–126. doi 10.1016/j.ajhg.2015.11.020

23. Rincent, R., D. Laloe, S. Nicolas, T. Altmann, D. Brunel, P. Revilla, V., M Rodriguez J. Moreno-Gonzalez, A. Melchinger, E. Bauer, C.C. Schoen, N. Meyer, C. Giauffret, C. Bauland, P. Jamin, J. Laborde, H. Monod, P. Flament, A. Charcosset, and L. Moreau. 2012. Maximizing the reliability of genomic selection by optimizing the calibration set of reference individuals: Comparison of methods in two diverse groups of maize inbreds (Zea may L.) Genetics 192:715–728 doi:10.1534/genetics.112.141473

24. De los Campos, G., Hickey, J. M., Pong-Wong, R., Daetwyler, H. D., & Calus, M. P. L. (2013). Whole-Genome Regression and Prediction Methods Applied to Plant and Animal Breeding. Genetics, 193(2), 327–345. http://doi.org/10.1534/genetics.112.143313

25. Jiang, Y., & Reif, J. C. (2015). Modeling Epistasis in Genomic Selection. Genetics, 201(2), 759–768. http://doi.org/10.1534/genetics.115.177907

26. Friedman, J., Hastie, T. and Tibshirani, R. (2008) Regularization Paths for Generalized Linear Models via Coordinate Descent, https://web.stanford.edu/~hastie/Papers/glmnet.pdf. Journal of Statistical Software, Vol. 33(1), 1–22 Feb 2010

27. Zou H, Hastie T. Regularization and variable selection via the elastic net. J. Royal. Stat. Soc. B. 2005;67(2):301–320. [

28. Park, T., Casella, G. (2008). The bayesian lasso. Journal of the American Statistical Association. 2008;103:681–686. 103.

29. Habier, DRL Fernando RL, Kizilkaya K and Garrick DJ(2011) Extension of the bayesian alphabet for genomic selection. BMC Bioinformatics 2011. 12:186.

30. Gianola, D., & van Kaam, J. B. C. H. M. (2008). Reproducing Kernel Hilbert Spaces Regression Methods for Genomic Assisted Prediction of Quantitative Traits. Genetics, 178(4), 2289–2303. http://doi.org/10.1534/genetics.107.084285

31. De los Campos, G., D. Gianola, G. J. M., Rosa, K. A., Weigel, and J. Crossa. 2010. Semi-parametric genomic-enabled prediction of genetic values using reproducing kernel Hilbert spaces methods. Genetics Research92 :295–308

32. Breiman L. “Random Forests”. Machine Learning 2001;45 (1): 5–32.

33. Gianola D, Okut H, Weigel KA, Rosa GJ. Predicting complex quantitative traits with Bayesian neural networks: a case study with Jersey cows and wheat. BMC Genetics. 2011;12:87. doi:10.1186/1471-2156-12-87.

34. Oury, F.X., Heumez, E., Rolland, B., Auzanneau, J., Bérard, P., Brancourt-Hulmel, M., Charrier, X., Chiron, H., Depatureaux, C., Falchetto, L., et al (2015) Winter wheat (Triticum aestivum L) phenotypic data from the multiannual, multilocal field trials of the INRA Small Grain Cereals Network. doi: 10.15454/1.4489666216568333E12

35. Rimbert H, Darrier B, Navarro J, et al. High throughput SNP discovery and genotyping in hexaploid wheat. Zhang A, ed. PLoS ONE. 2018;13(1):e0186329. doi:10.1371/journal.pone.0186329.

36. Zhou Y, Vales MI, Wang A, Zhang Z. Systematic bias of correlation coefficient may explain negative accuracy of genomic prediction. Brief Bioinform. 2016:bbw065 doi:10.1093/bib/bbw064

37. Endelman, J.R., and J.L. Janninck. Shrinkage estimation of the realized relationship matrix. G3: Genes, Genomes, Genetics 2012, 2:1405–1413 doi: 10.1534/g3.112.004259

38. Wimmer, V., Albrecht, T., Auinger, J.J., Schön; C.C. 2012. Synbreed: a framework for the analysis of genomic prediction data using R, Bioinformatics, 28: 2086–2087

39. Ma, C., Cheng, Q., Qiu Z., Song, J. (2017). Package ‘G2P’ Genomic selection Prediction and Evaluation https://github.com/cma2015/G2P

40. Heslot, N., Yang, H.P., Sorrells M.E., Jannink, J.L. 2012. Genomic Selection in Plant Breeding: A Comparison of Models. Crop Sci. 52:146–160

41. Michel, S., Ametz, C., Gungor, H., Epure, D., Grausgruber, H., Löschenberger, F., & Buerstmayr, H. (2016). Genomic selection across multiple breeding cycles in applied bread wheat breeding. TAG. Theoretical and Applied Genetics. Theoretische Und Angewandte Genetik, 129, 1179–1189. http://doi.org/10.1007/s00122-016-2694-2

42. He, S., Schulthess, A.W., Mirdita, V., Zhao, Y., Korzun, V., Bothe, R., Jochen, E. and C. Reif (2016) Genomic selection in a commercial winter wheat populational. Theor Appl Genet (2016) 129: 641. https://doi.org/10.1007/s00122-015-2655-1

43. Liu, X., Wang, H., Wang, H., Guo, Z., Xu, X., Liu, J., Wang, S., Li, W.X., Zou, C., Prasanna, B.M., Olsen, M.S., Huang, C., Xu, Y. (2018), Factors affecting genomic selection revealed by empirical evidence in maize, The Crop Journal. https://doi.org/10.1016/j.cj.

44. Ly, C., Toghiani, S., Ling, A.,, Aggrey, S.E. 3,4, Rekaya, R. (2018). High density marker panels, SNPs prioritizing and accuracy of genomic selection. BMC Genet. 2018 Jan 5;19(1):4. doi: 10.1186/s12863-017-0595-2.

45. Heffner E.L., Jannink J-L. Sorrells M.E (2011) Genomic Selection Accuracy using Multifamily Prediction Models in a Wheat Breeding Program. The Plant Genome 4:65–75.

46. Ma, Y., Reif, J.C., Jiang, Y. et al. Potential of marker selection to increase prediction accuracy of genomic selection in soybean (*Glycine max* L.) Mol Breeding (2016) 36: 113. https://doi.org/10.1007/s11032-016-0504-9

47. Cericola C, Jahoor A, Orabi J, Andersen JR, Janss LL, Jensen J. Optimizing training population size and genotyping strategy for genomic prediction using association study results and pedigree information. A case study in advanced wheat breeding lines. PLoS ONE (2017);12 (1) e0169606. Doi 10.371/journal.pone.0169606

48. Schulz-Streeck, T., Ogutu, J. O., & Piepho, H.-P. (2011). Pre-selection of markers for genomic selection. BMC Proceedings, 5(Suppl 3), S12. http://doi.org/10.1186/1753-6561-5-S3-S12

49. Daetwyler, H.D., B. Villanueva, and J.A. Woolliams. 2008. Accuracy of predicting the genetic risk of disease using a genome-wide approach. PLoS ONE 3:e3395. doi:10.1371/journal.pone.0003395

50. Daetwyler, H.D., R. Pong-Wong, B. Villanueva, and J.A. Woolliams. 2010. The impact of genetic architecture on genome-wide evaluation methods. Genetics 185:1021–1031. doi:10.1534/genetics.110.116855

51. Daetwyler, H.D., Calus, M.P.L., Poing-Wong, R., De Los Campos, G., Hickey, J.M. 2013. Genomic Prediction in Animals and Plants: Simulation of Data, Validation, Reporting, and Benchmarking. Genetics 193: 347–365

52. Isidro, J., Jannink, JL., Akdemir, D., Poland J., Heslot, N., Sorrels, M.E. (2015) Training set optimization under population structure in genomic selection. Theor Appl Genet (2015) 128: 145. https://doi.org/10.1007/s00122-014-2418-4

53. Rincent, R., Charcosset, A. & Moreau, L. Predicting genomic selection efficiency to optimize calibration set and to assess prediction accuracy in highly structured populations. Theor Appl Genet (130: 2231. https://doi.org/10.1007/s00122-017-2956-7

54. Akdemir D, Sanchez JI, Janninck JL.(2015) Optimization of genomic selection training populations with a genetic algorithm. Genetics Selection Evolution 2015;47: 38. Doi 10.1186/s12711-015-0116-6

55. Song, J., Carver, B.F., Powers, C, Yan, L., Klapste, J., El-Kassaby, Y.A., Chen, C. (2017) Practical application of genomic selection in a doubled-haploid winter wheat breeding programme. Mol Breeding (2017) 37: 117. https://doi.org/10.1007/s11032-017-0715-8

56. Wang, X., Xu, Y., Hu, Z., Xu, C. (2018), Genomic selection methods for crop improvement: Current status and prospects. The Crop Journal (2018) https://doi.org/10.1016:j.cj/2018.03.001

57. Wang, D., El-Basyoni, I.S., Baenziger, P.S. Crossa, J., Eskridge, K.M., Dweikat I. (2012) Prediction of genetic values of quantitative traits with epistatic effects in plant breeding populations. Heredity, 109 (2012), pp. 313–319

58. González-Camacho, J. M., de los Campos, G., Pérez, P., Gianola, D., Cairns, J. E., Mahuku, G., Crossa, J. (2012). Genome-enabled prediction of genetic values using radial basis function neural networks. TAG. Theoretical and Applied Genetics., 125(4), 759–771. http://doi.org/10.1007/s00122-012-1868-9

59. Li B, Zhang N, Wang Y-G, George AW, Reverter A and Li Y (2018) GenomicPrediction of Breeding Values Using aSubset of SNPs Identified by ThreeMachine Learning Methods.Front. Genet. 9:237.doi: 10.3389/fgene.2018.00237

60. Ma, W., Qiu, Z., Song, J., Li, J., ., Cheng, Q., Zhai, J., Ma C(2018) Deep convolutional neural network approach for predicting phenotypes from genotypesPlanta (2018) 248: 1307. https://doi.org/10.1007/s00425-018-2976-9

61. Kang, H., Zhou, L., Liu, J. (2017) Statistical considerations for genomic selection. Front. Agr. Sci. Eng. 2017, 4(3):268–278.

62. Spindel, J.E., Mc Couch, S.R. (2016) When more is better: how data sharing would accelerate genomic selection of crop plants. New Phytologist (2016) 212:814–826

63. Heslot, N., Jannink, J.L., Sorrells M.E. (2015). Perspectives for Genomic Selection. Application and Research in Plants. Crop Sci. 55:1–12

64. Bassi FM, Bentley AR, CharmetG, Ortiz R, Crossa J. Breeding schemes for the implementation of genomic selection in wheat (Triticum spp.). Plant Science 2016;242:23–36. Doi10.1016/j.plantsci.2015.08.021

65. J. Crossa, P. Pérez-Rodríguez, J. Cuevas, O. Montesinos-López, D. Jarquín, G. delos Campos, J. Burgueño, J.M. Camacho-González, S. Pérez-Elizalde, Y. Beyene, S. Dreisigacker, R. Singh, X. Zhang, M. Gowda, M. Roorkiwal, J. Rutkoski, R. Varshney. Genomic selection in plant breeding: methods, models, and perspectives. Trends Plant Sci., 22 (2017), pp. 961–975, 10.1016/j.tplants.2017.08.011

66. Michel, S., Kummer, C., Gallee, M., Hellinger, J., Ametz, C., Akgöl, B., Epure, D., Loschenberger, F., Buerstmayr, H. (2018). Improving the baking quality of bread wheat by genomic selection in early generations. TAG. Theor Appl Gen. 131(2), 477–493. http://doi.org/10.1007/s00122-017-2998-x

